# Classification and Analysis of Minimally-Processed Data from a Large Magnetoencephalography Dataset using Convolutional Neural Networks

**DOI:** 10.1101/846964

**Authors:** Jon Garry, Thomas Trappenberg, Steven Beyea, Timothy Bardouille

## Abstract

Convolutional neural networks were used to classify and analyse a large magnetoencephalography (MEG) dataset. Networks were trained to classify between active and baseline intervals of minimally-processed data recorded during cued button pressing. There were two primary objectives for this study: (1) develop networks that can effectively classify MEG data, and (2) identify the important data features that inform classification. Networks with a simple architecture were trained using sensor and source-localised data. Networks trained with sensor data were also trained using varying amounts of data. The important features within the data were identified via saliency and occlusion mapping. An ensemble of networks trained using sensor data performed best (average test accuracy 0.974 ± 0.001). A dataset containing on the order of hundreds of participants was required for optimal performance of this network with these data. Visualisation maps highlighted features known to occur during neuromagnetic recordings of cued button pressing.

## 1. Introduction

Big data in medical imaging has the potential to inform new models in diagnostics. In recent years there has been an exponential increase in the amount of data in many fields, including health care [1]. For example, the Cambridge Centre for Ageing and Neuroscience (Cam-CAN) has provided a large open-access dataset that includes demographic, behavioural, and structural and functional neuroimaging data recorded across a large, healthy adult population, for the purpose of investigating healthy ageing in a generalizable manner [2]. Of particular relevance for this study, the Cam-CAN dataset includes magnetoencephalography (MEG) recordings during a simple cued button press task and structural magnetic resonance images (MRIs) in the same participants. With these appropriately large datasets, approaches found in the field of deep learning can be applied in order to augment and improve currently established methods for statistical analysis in neuroimaging, with the end goal of improved understanding of the human brain and potentially better diagnosis for patients [3].

Whereas traditional machine learning uses models that act as discriminators in classifying or clustering data, deep learning models generate latent representations of input data. This is referred to as representation learning and it is central to deep learning [4]. One common model in deep learning is the convolutional neural network (CNN), which is a type of artificial neural network originally designed for processing images and video [5, 6]. A CNN contains layers that convolve inputs with small matrices known as kernels in order to extract specific types of information. Kernel weights are adjusted during training, along with neuron connections, to improve classification accuracy. When many convolutional layers are stacked together, a nested hierarchy is formed allowing for the detection of complex structure within the data [4]. A CNN therefore performs a combination of efficient feature extraction (via weight sharing) and functional approximation.

Neural networks had been criticised for being difficult to interpret when compared to models developed using more traditional machine learning techniques. However, researchers in computer vision and image recognition have developed methods for investigating and visualising trained networks. Some of these methods include activation mapping [7, 8], saliency mapping [7], and occlusion mapping [9], which can identify the portions of an input that contribute most to strongly successful classification. Combining deep learning, large functional neuroimaging datasets, and formal visualisation techniques opens new avenues for characterising which brain signals are most salient, towards potentially identifying task-specific or pathological activity. As an initial step in this direction, we are interested in developing models of healthy brain function constructed using large collections of normative data with a simple, well-understood task. In this validation framework, we can determine if the data representations learned by a CNN for a large MEG dataset acquired during cued button pressing will match expectations based on previous literature.

The MEG correlates of cued-button pressing have been well-studied. Auditory and visual cues both generate a reproducible complex of magnetic field deflections over a time period of approximately 300 ms following the cue. The deflections, termed the auditory and visual evoked fields (AEF and VEF, respectively) [10], are observed bilaterally on sensors over the temporal and occipital cortex, respectively. As well, the suppression of “alpha” (~10 Hz) bursts is observed on occipital sensors for up to one second when a visual stimulus occurs [11]. The button press is associated with a reproducible complex of magnetic field deflections that includes pre- and post-movement components observed on sensors over the primary motor cortex (M1), the primary somatosensory cortex (S1) and the broader sensorimotor network [12]. As well, centrally-generated “mu” (~8-12 Hz) and “beta” (~15-30 Hz) bursts are suppressed for about 750 ms following a button press and enhanced 750 ms to 2000 ms post-movement (i.e., post-movement beta rebound) [11, 13]. When a simple button press is cued by auditory and visual stimulus, we expect that all of these responses will summate, with little change to the spatiotemporal dynamics.

In this paper, a CNN is presented which is designed to classify between minimally-processed MEG measurements recorded during active and baseline intervals of a cued button pressing task in the Cam-CAN large dataset. Although MEG measurements typically have a low signal-to-noise ratio, there is underlying structure that is consistent across sets of channels and times which a CNN can learn. It is hypothesised that, with enough training data, a CNN will learn to effectively classify MEG records. The performance of networks that have been trained using minimally processed sensor records and source estimated data are compared. Visualisation methods are presented in order to reveal the signals that these networks are extracting for accurate classification. It is hypothesised that the visualisation of the CNN will reveal a sensitivity to occipital and temporal activity (due to the cue) and motor-related activity (due to the button press).

## 2. Methods

### 2.1. Participants & Paradigm

Data used in the preparation of this work were obtained from the Cam-CAN repository (available at http://www.mrc-cbu.cam.ac.uk/datasets/camcan/). Data were recorded during the second stage of the Cam-CAN study [2], which included 700 individuals equally distributed across sex and age between 18 and 87 years of age. MEG data were acquired from 306 channels (Elekta Neuromag, Helsinki, Finland) at a sampling rate of 1000 Hz with an inline band-pass filtering between 0.03 and 330 Hz using a 306-channel Vectorview system. Digitisation of anatomical landmarks and additional points on the scalp was also performed for registration of MEG with MRI. Head position was monitored continuously, and electrooculogram and electrocardiogram were recorded concurrently along with stimulus/response event markers. During the MEG recording, participants pressed a button using their right index finger after the presentation of visual, auditory, or combined audio-visual stimuli. The inter-stimulus interval was randomised between two and 26 seconds in order to prevent anticipatory effects, and participants performed 128 trials. T1-weighted magnetic resonance images (MRI) were acquired using the 3T Siemens Tim Trio system with a 32-channel head coil.

### 2.2. Data Preparation

Data pre-processing performed by the Cam-CAN group included the application of temporal signal space separation (tSSS) [14] to remove environmental magnetic noise, perform head movement corrections, and transform data to a common head position [15]. As previously reported [16], in-house pre-processing included splitting up each participant’s dataset into 3.4 s trials centered on each button press and independent component analysis (ICA) [17] for artefact removal. Trials were excluded for poor task performance (response time greater than one second) or if the button press occurred within three seconds of the previous button press (contaminated baseline). The number of features were reduced by including only the 102 magnetometer channels, and excluding the planar gradiometers. Both sensor types are sensitive to cue- and response-related activity, and including only one sensor orientation simplified visualisation (as opposed to two planar orientations). Since we were interested in evoked field and rhythms below approximately 30 Hz, a low-pass filter of 40 Hz was applied to all trials and data were down-sampled to 250 Hz. For each participant, these processes resulted in MEG trial data as a tensor with dimensions [# Trials, # Sensors, # Time Samples]. Importantly, the resultant data has undergone minimal processing for input to the CNN, specifically to test if the network will learn representations that match previously reported task-related activity. As well, MEG trial data were averaged across all trials and participants to generate the grand-average evoked field data for cued button pressing. These data were visualised to reveal the average magnetic field deflections associated with the task.

For input to the CNN, one-second MEG data segments centred on the button press were used as a record belonging to the “Active” class. Given that participant response times were 300 ms on average [16], the active records included both cue- and motor-related activity. One-second MEG data segments ending 700 ms prior to the button press were extracted from each trial as “Baseline” class records. Thus, each participant trial provided two classified records. All records were scaled to unit normal by subtracting the mean and dividing by the standard deviation across each sensor. After all processing tasks were complete, there were 75,396 records across 605 participants with an even distribution between the active and baseline classes. Each record was represented as an image with each row representing one of the 102 MEG sensors and each column representing one of the 250 time samples.

Classification was also performed on representations of the sensor data transformed into source space providing more spatially-specific data. Source estimation was performed by applying dynamical Statistical Parametric Mapping (dSPM) [18] to the MEG trial data. All sensors (magnetometers and gradiometers) were kept in order to provide more accurate source estimation. A boundary element model based upon each participant’s MRI was used to provide an accurate model for source localisation, with MEG to MRI co-registration performed manually [16]. Time courses of source estimates were generated at the centre of mass of each of 68 anatomically-defined regions of interest (FreeSurfer aparc cortical parcellation) [19]. Source estimated trials were split to form records and normalised in the same manner used for sensor records. Thus, each record consisted of 68 anatomical regions by 250 time samples. There were a total of 72,518 source-level records from 582 participants. Twenty three participants were excluded due to challenges in the source estimation process, such as missing data or errors in MR reconstruction and registration.

### 2.3. Deep Learning

For classification (sensor and source level), the records were split into training (80%), validation(5%), and testing (15%) subsets, with pseudorandom sampling such that all records from a specific participant only existed within a single subset to avoid data leakage. The CNN consisted of four layers: two convolutional layers, one fully-connected layer, and a softmax classification layer. The first convolutional layer contained eight 8 x 16 (channels x time samples) kernels that were convolved with the input MEG records (zero padding with a stride of one) to produce eight feature maps. This kernel size, as opposed to the standard 3 x 3 kernel, was chosen in order to capture structure on the timescale of tens of milliseconds, and because MEG data is spatially less resolved than image data. The second layer performed convolutions using sixteen 3 x 3 kernels. A standard kernel dimension was used in the second layer because we had no *a priori* justification for a different kernel dimensionality. The outputs were flattened and connected to a fully-connected layer, which contained 64 neurons for sensor records and 32 neurons for source-estimated records. Each layer used standard rectified linear units (ReLU) for non-linear activation. Finally, a two neuron layer with softmax activation transformed the two class neuron values to output a probability distribution across the active and baseline classes.

Batch normalisation was employed within each layer prior to passing the weighted sums to the ReLU activation functions to provide gradient stability between layers, and to speed up network training [20]. In order to prevent over-fitting to the data, the network used dropout regularisation in which a proportion of network weights were randomly excluded during each training step. The fully-connected layer excluded 50% of neurons during each training step, and the convolutional layers excluded 25% of the feature maps at each step.

The AdaGrad algorithm was used to minimise binary cross-entropy during training [21]. Batches of 25 records randomly sampled from the training set were used at each training step when using the sensor records, whereas networks trained on the source-estimated records used batches of 50 records. A training epoch was complete when all of the available training data had been used to train the network. After each epoch, the classification accuracy of the network was calculated using the training and validation datasets. The process of training the networks was repeated for 50 epochs. Optimal networks were then chosen based upon a combination of largest validation accuracy achieved and smallest associated cross-entropy values. Ensembles of 10 identical networks were trained using all of the available data in order to characterise the performance of our network design.

Initially, one ensemble was trained using the sensor record data and a second ensemble was trained with the source localised data. Due to the poor performance of networks trained using source localised records (see Results), further investigations were limited to the sensor data only. To characterise network performance in terms of dataset size, sub-datasets with varying numbers of participants were constructed by randomly sampling participants from the original sensor records, and repeating the deep learning process. For training on sub-datasets, ensembles of 100 identical networks were trained using each dataset for a maximum of 15 epochs. Due to the initially smaller dataset sizes and a constant model capacity, a higher degree of variability in performance was expected due to over-fitting. As such, a larger number of networks were included in each ensemble to capture this variability. Training was limited to 15 epochs in order to limit the degree to which the networks would over-fit.

### 2.4. Network Visualisation

For the purposes of visualisation and attribution analysis, a single network was selected which was fully trained upon MEG sensor records. From the ensemble of 10 identically initialised and trained networks described above, we selected the network that exhibited the best performance in terms of classification accuracy and cross-entropy loss from the ensemble. Visualisation methods were then performed using this network in order to investigate the features specific to MEG that informed model performance.

The eight first-layer kernels were convolved with all 11,396 records in the test dataset in order to generate eight sets of feature maps. To examine sensitivity to so-called “evoked” field deflections, inter-record averages were computed across active class records for each of the eight sets of feature maps. These maps were compared with the grand-average MEG data. Additionally, we investigated feature map sensitivity via peak detection in each map (rather than the grand-average). Positive and negative peaks with magnitude greater than or equal to four standard deviations in each record were identified and the peak channel and time was recorded across all feature maps for each of the eight kernels. From these data, two-dimensional histograms (102 sensors x 250 time samples) that counted the number of times a peak was detected for a particular sensor and time were generated for each kernel to investigate which signal components were commonly accentuated during propagation through the first convolutional layer.

Saliency maps provide a representation of which individual data points within a given record have the largest impact on classification [7]. In order to generate these maps, the softmax activation function was removed and the values of the class-specific neurons were computed directly upon processing input. The saliency of each data point was mapped by calculating the gradient of the score function with respect to each data point within an input record. In order to investigate the most commonly salient points in our input records, saliency maps were generated across all records within the test dataset, and peak analysis (as described above) was also performed on the saliency maps. However, since saliency is positive-definite, only the positive peaks outside of four standard deviations were used.

Occlusion maps were generated by systematically setting portions of the input record to zero and recording the change in the score function for all test records. A 2 x 2 (channels x time samples) occlusion window was used, and the resulting values were re-scaled to the range [-0.5,0.5] for each record. Thus, those regions most important for classification provided the largest negative values within the occlusion map. The channels and times that contained negative peaks outside of four standard deviations were recorded and visualised via 2-D histogram.

## 3. Results

The networks trained on the MEG sensor data converged to an average validation accuracy of 0.960 ± 0.001 after six epochs, with the average loss function attaining a minimum around the same epoch before trending towards an increase. The variance in the validation cross-entropy trended towards an increase after epoch six, suggesting that networks tended to over-fit to the data after this point. The average classification accuracy calculated over the test dataset and using optimal network configurations was 0.974 ± 0.001. There was a relatively even split in mislabelled records between the active and baseline classes. The networks trained on the source estimated data did not perform as well as those trained on the sensor data. The average validation accuracy reached a maximum of 0.814 ± 0.009 after four epochs, and the classification accuracy on the test subset was 0.793 ± 0.006. Due to the poor performance of the CNN with source estimated data, further analysis focused on deep learning with the sensor data. Thus, all results that follow relate to the sensor data only.

Figure 1 shows the dependence of network performance on the number of participants included within the dataset. From the smallest dataset containing 20 participants up to the dataset containing 200 participants, classification accuracy steadily increased suggesting that a substantially better model could be developed as more participants were included in the analysis. For datasets larger than 200, the networks tended to increase in accuracy but with a smaller rate of increase suggesting that adding participants does not substantially impact model performance at these samples sizes. The average test accuracy calculated over the ensemble trained using data from 600 participants was 0.965 ± 0.002.

**Figure 1:**
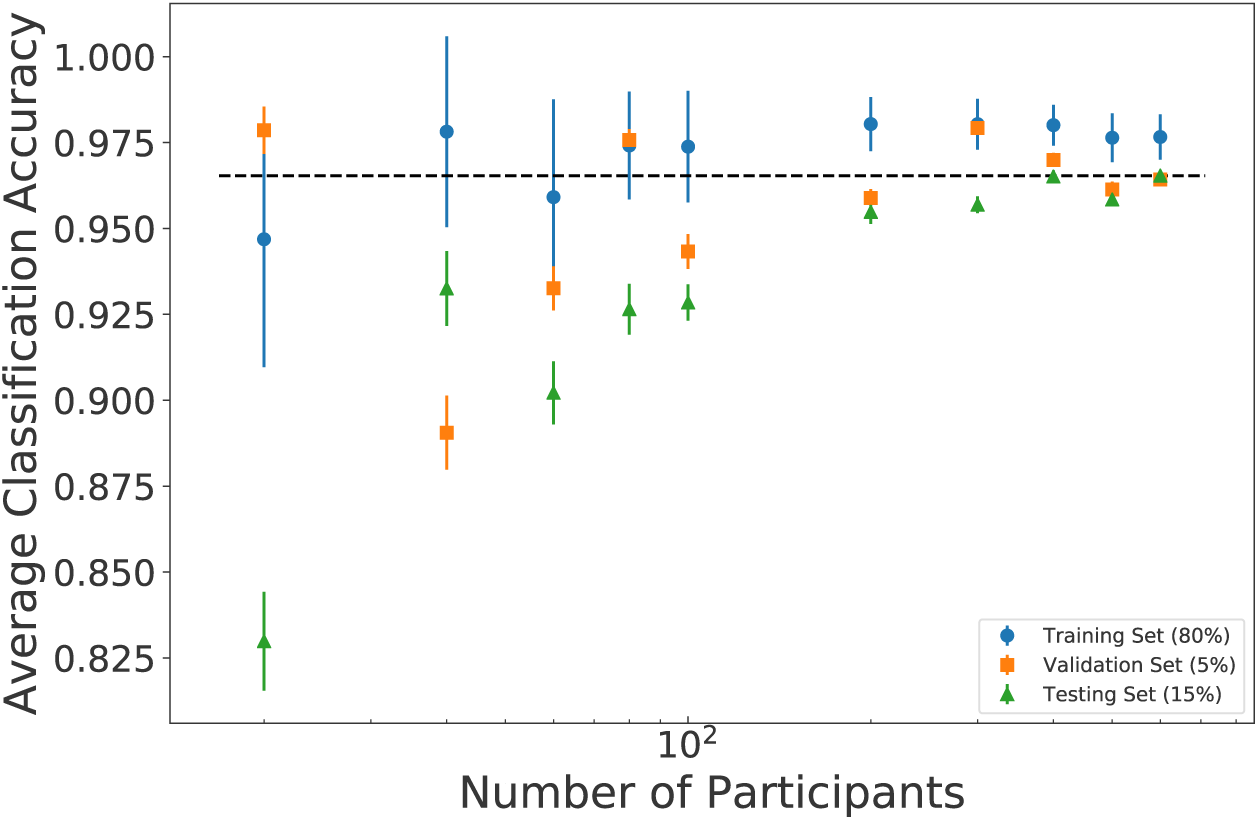
Network performance as a function of dataset size. Each point represents the average classification accuracy for the training set (blue circles), validation set (orange squares), and testing set (green triangles). The dashed line indicates average test accuracy when the network was trained on 600 participants. Accuracy steadily increases up to datasets containing 200 participants, then increases at a lower rate.

Figure 2 shows the grand-average MEG data for the 500 ms prior to and following the button press. Clear magnetic field deflections are evident at −100 ms, −60 ms, 55 ms, 120 ms, and 325 ms. The topography of the peaks prior to the button press suggest likely bilateral occipital activity related to the cue. At 55 ms, the topography suggests bilateral central sources, likely related to efferent and afferent sensorimotor activity associated with button pressing. The topographies at 120 ms and 325 ms are less easily interpreted, likely due to the involvement of multiple brain regions in processes occurring at these latencies.

**Figure 2:**
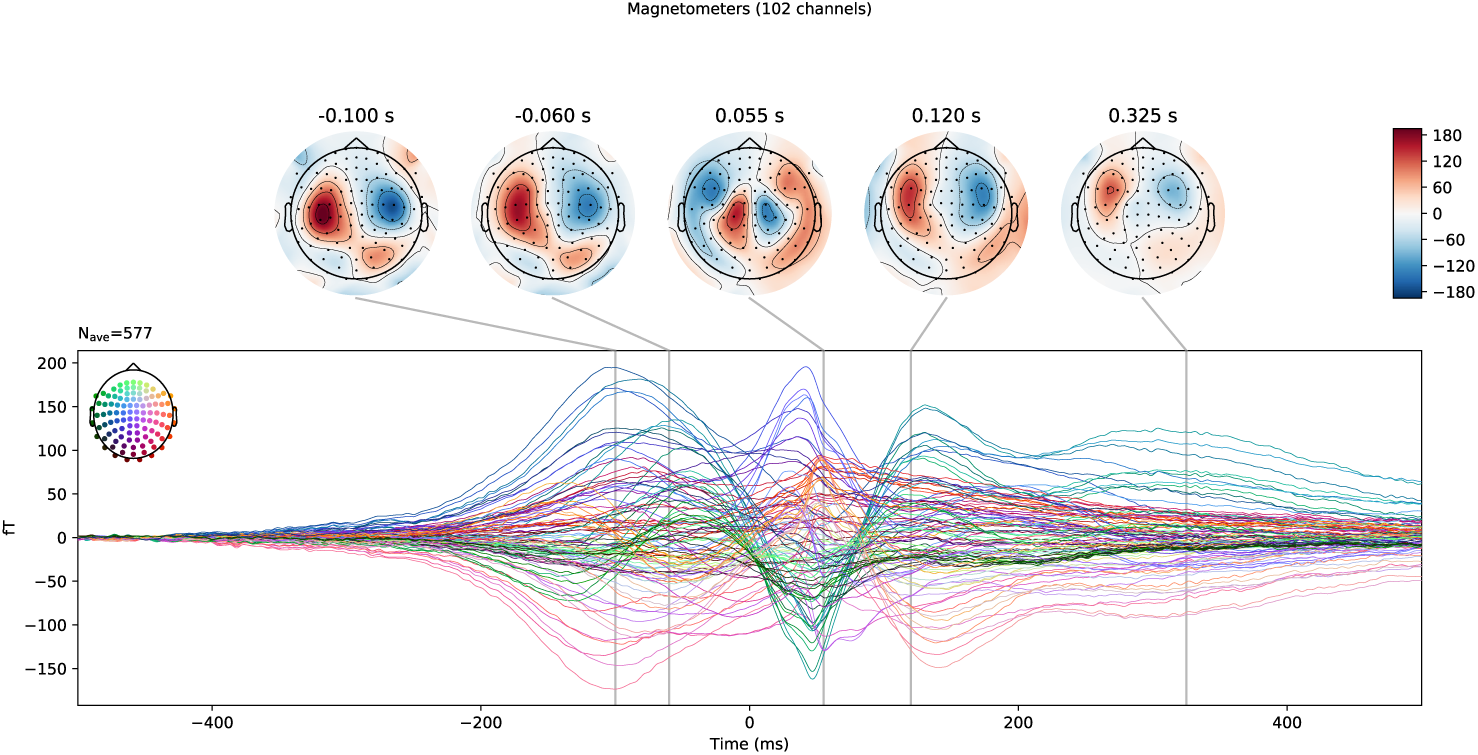
The grand-average MEG data for 500 ms before and after the button press. Each line on the graph represents average magnetic field deflections for a single MEG sensor. The topographic plots are shown above for the peak latencies of evident field deflections. The topographies suggest cue-related bilateral occipital activity, and response-related sensorimotor activity.

Figure 3 compares the grand-average of the active test records to the grand-average of the feature maps generated using a single first layer kernel. The grand-average feature map shows structure similar to that of the grand-average of unconvolved test records, indicating that the kernel is sensitive to the evoked fields occurring during a cued button-press task. The peaks within the active feature maps tended to have larger amplitudes over a different set of sensors compared to the grand-average, indicating differences in spatial sensitivity. Furthermore, some of these peaks were temporally shifted as compared to the associated grand-average peaks, indicating that the kernel is possibly sensitive to some derivative of the temporal dynamics.

**Figure 3:**
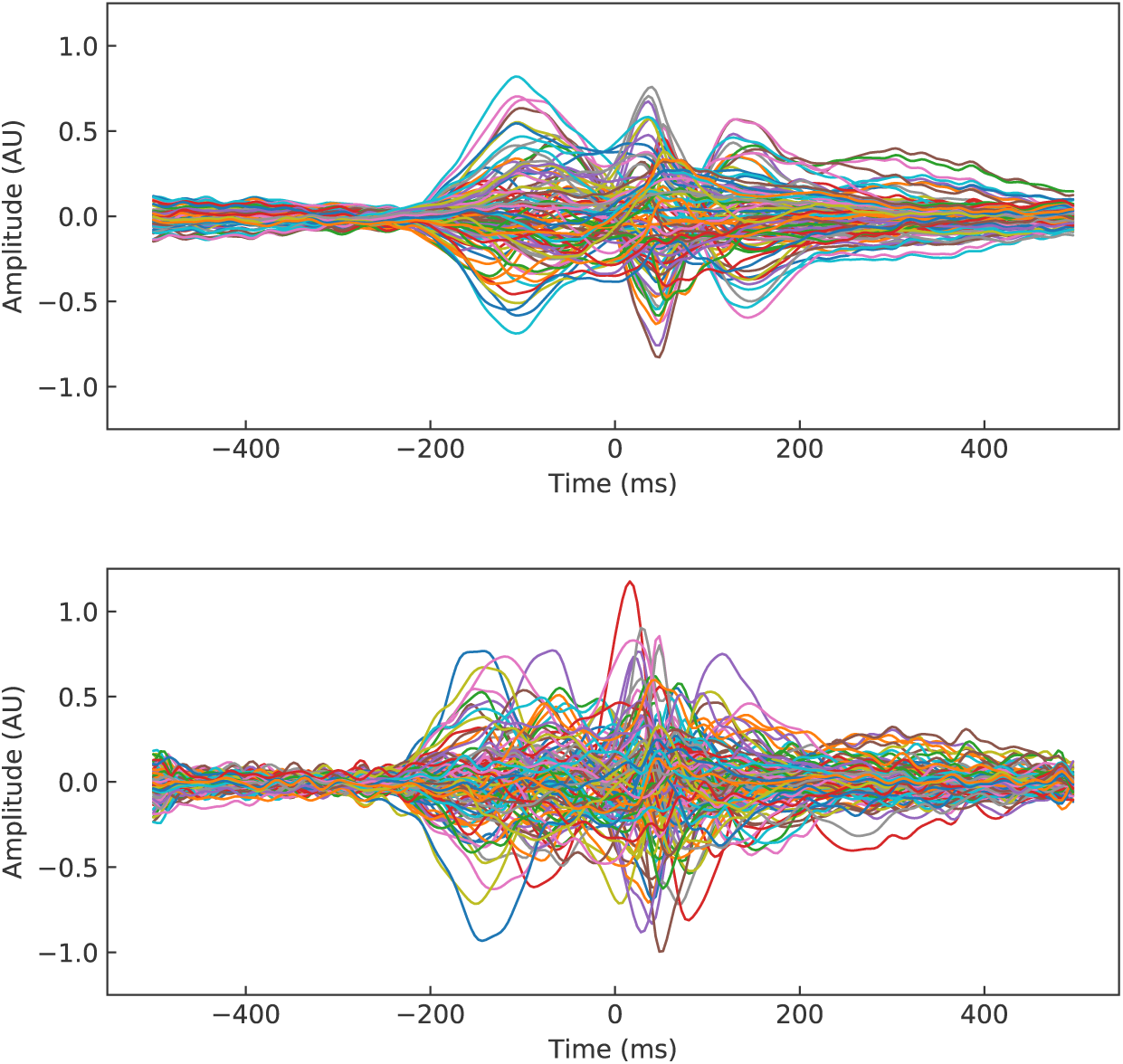
A comparison of the grand-averaged calculated test records (top) and feature maps generated from a first layer kernel (bottom). Each line represented data from one row of the grand-averaged record. Plots are centred at the button press at time *t* = 0 ms. Similar peaks, associated the cue and button press, can be seen within both averages. Feature map peaks have different relative amplitudes and timing with respect to their grand-average counterparts, indicating some shift in sensitivity following convolution.

Figure 4 shows a 2-D histogram of peaks in the feature maps produced by the first layer kernel, to reveal when and where the largest signals tended to exist. Two peaks were observed in the feature map histogram: one prior to button press between −200 ms to 0 ms, and one afterwards between 0 ms to 150ms. These are likely associated with expected cue- and response-related peaks, respectively. A key feature observed within this (and other) histograms was an apparent decrease in counts after the button press, in comparison to the interval prior to the button press. This decrease in counts matches the temporal dynamics of beta suppression in S1/M1, which occurs following movement onset. The topographic map constructed by mapping the decrease in peak counts following the button press is shown on the right side of Figure 4. The largest decreases in counts occurs over the central sensors overlying S1/M1, as would be expected with mu or beta suppression.

**Figure 4:**
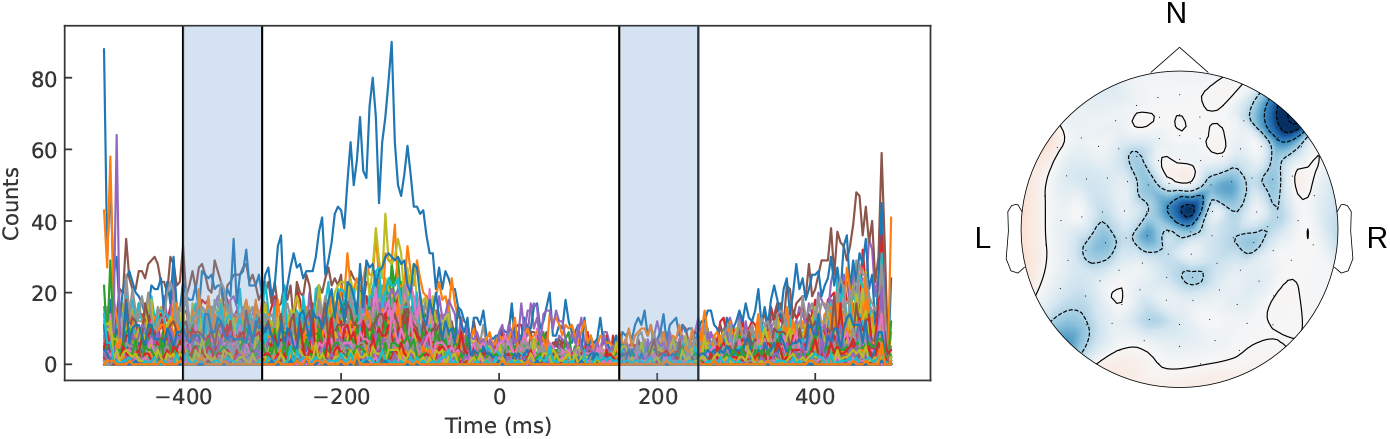
Peak analysis of the feature maps show evidence that trained kernels were sensitive to beta rhythms. A histogram of peaks in feature maps for one kernel (left) is shown with each line representing counts for one row in the feature map. A decrease in counts following the button press between the shaded areas is observed. This difference in counts between shaded intervals was plotted topographically (right), indicating a central generator, in keeping with mu or beta suppression.

Along with cortical rhythm suppression, this particular kernel appeared to be sensitive to features along the edge of sensor space. This could be the result of the representation used for the sensor records. Within each record, sensors were listed by row in an order that matched the naming convention set by the hardware manufacturer. This ordering only partially resembled the spatial relationship among the sensors with groups of sensors clustered together within specific regions of the head. Importantly, there were some spatial discontinuities in the sensor list. Possibly, these spatial discontinuities in the MEG data were perceived as large edges in feature maps, leading to a high number of peak counts at the edge of the sensor space.

Figure 5 shows the peak histograms for saliency maps (top) and occlusion maps (bottom) generated across the active and baseline records (left to right). Within the saliency map histograms, the distribution of peaks is similar across both classes with a large peak occurring between *t* = 50 ms and *t* = 60 ms on channels below channel 50. Smaller peaks precede and follow the large peak on a subset of the same channels. There are also clusters of peaks prior to button press on channels 60 and higher. The structure contained within these histograms suggest that the network was sensitive to activity likely related to both the cue (*t* < 0 ms) and the button press (*t* > 0 ms). The similarity between the active and baseline histograms could possibly be due to the fact that the network attempted to identify the occurrence or absence of these features when performing classification. Some slight differences between active and baseline saliency is apparent, however. The active class appears to contain more salient data points prior to button press between *t* = −200 ms and *t* = 0 ms, as compared to the baseline class. Conversely, more salient points occurred after *t* = 200 ms in the baseline records, as compared to the active records. There are common structures when comparing the saliency and occlusion map histograms. Large central peaks can be observed between *t* = 50 ms to *t* = 60 ms within both the active and baseline histograms. Although there are similar peaks within both active and baseline histograms of the occlusion maps, the baseline counts are far more distributed over time, with relatively more peaks counted after *t* = 200 ms as shown in the marginal distribution for time.

**Figure 5:**
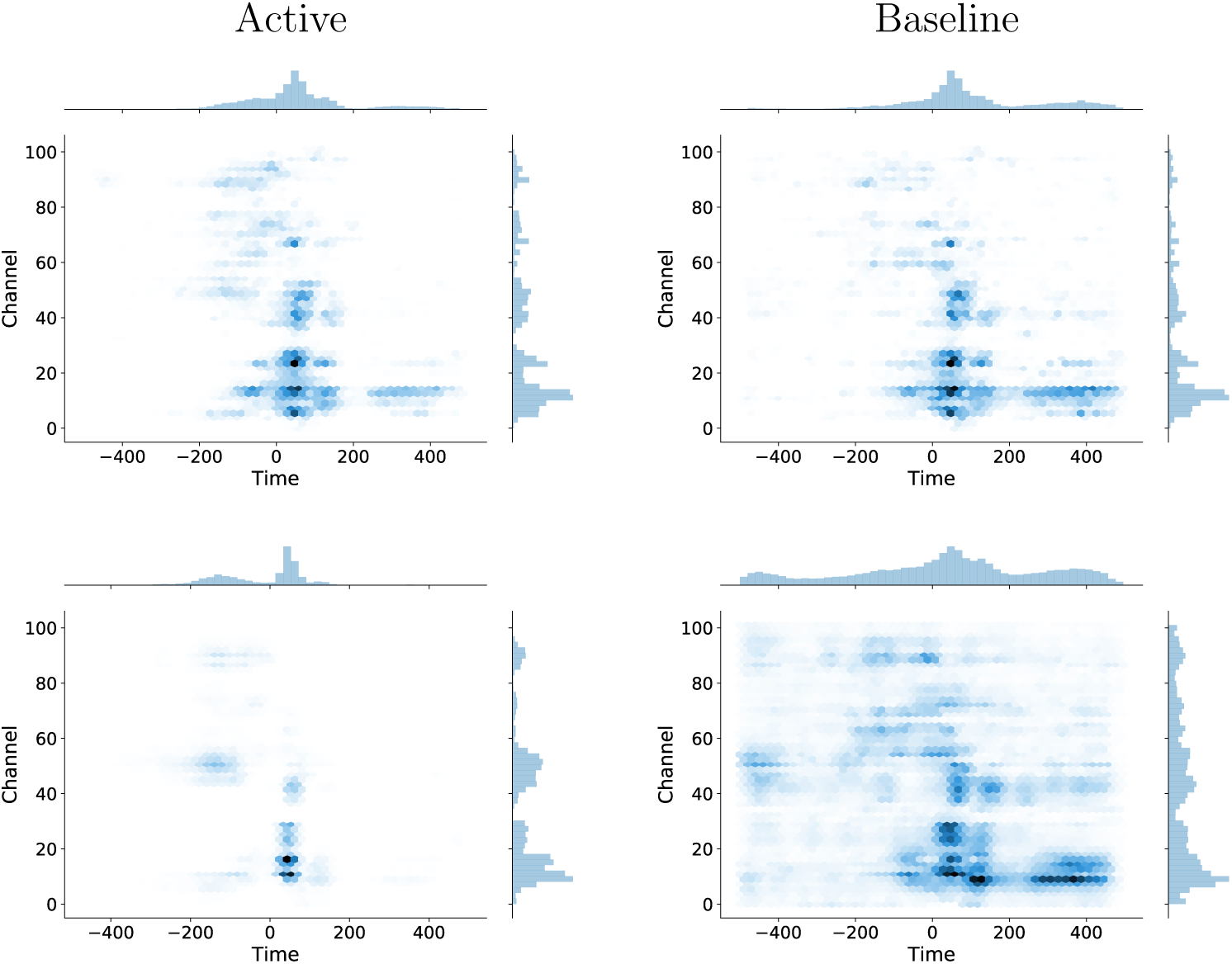
2-D histograms are shown that count peaks in saliency (top) and occlusion (bottom) for the active (left) and baseline (right) test records, indicating the importance of particular data points in classifying the input record.

Figure 6 shows topographic representations of the test records and visualisation maps over five selected time intervals for the active test records. As in Figure 2, the test record topographies suggests bilateral occipital activity related to the cue prior to button press, and bilateral activity related to the button press on central channels, with less obviously interpretable topographies occuring later. The topographic representation of peaks in the saliency maps appears to have a strong correspondence to the features observed in the grand-average of test records, as well as our expectations based on prior studies of cued-button pressing. Specifically, during −200 to 0 ms, important features in central and occipital sensors were observed that match closely with the sensors activated in the test records. During −65 to −55 ms, high counts on a small locus of left central sensors was observed, which overlaps with a positive locus in the test records. Interestingly, the other strongly activated sensor loci in the grand-average at this time interval show a weaker correspondence in the saliency map, indicating that not all of the field represented in the grand-average is consistently salient for classification by the CNN. During the remaining three time intervals, consistently high saliency is measured on central sensors that overlap to lesser degree with the grand-average test records. Interestingly, these later saliency topographies match more closely with the feature map topography in Figure 4. This points to a possible representation of cortical rhythm suppression in the saliency map.

**Figure 6:**
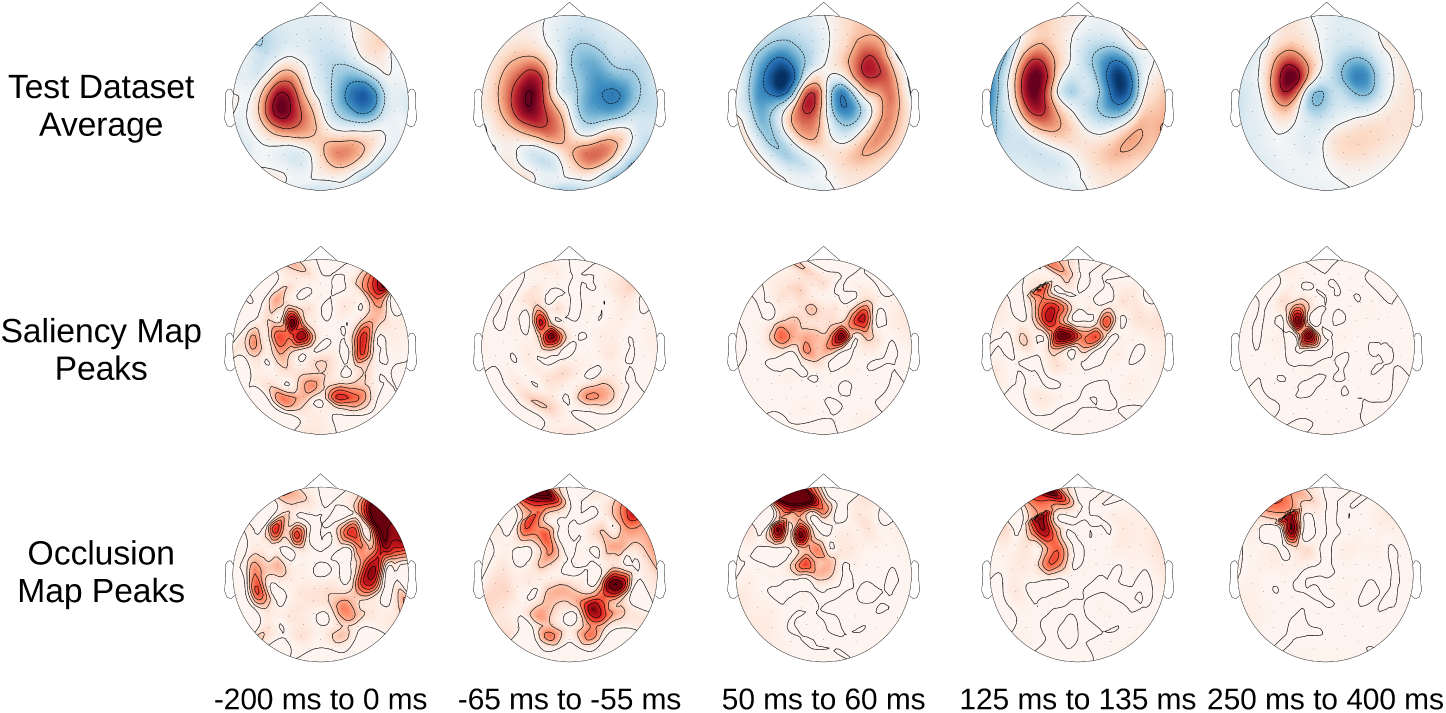
Topographic plots of the grand-average test records and visualisation histograms are shown for five time intervals of interest.

The topographic maps of occlusion data also identified some of the same regions of activity as saliency, including loci of sensors along the edge of the array. These edge effects are uncommon in results generated by standard approaches to analysing neuromagnetic recordings, and likely represent an artefact of the spatial discontinuities in the representation of the records, as previously described. Kernels that are selective for sensors (rather than time) may be more sensitive to these discontinuities in the data representation.

## 4. Discussion

In this study we developed a CNN that can classify between active and baseline intervals of minimally processed MEG data. We showed that high classification accuracy can be achieved using a relatively small CNN. Using visualisation techniques, we showed that the CNN learns the expected underlying structure within the data, suggesting that a CNN is a viable choice for constructing models to represent MEG data. Whereas the data used in this study contain activity from a well known task, this research paves the way for the development of networks that can identify activity in more complicated tasks and pathological activity within a clinical setting. We also showed that network performance increased dramatically with dataset size up to 200 participants, with less pronounced increases in classification accuracy for larger sample sizes. These results suggest that any future application of deep learning to minimally-processed MEG data should employ datasets containing on the order of hundreds of participants.

Although our CNN could effectively classify sensor-level records, networks trained on the source-localised data produced poor results. This may have occurred because of the spatial representation of the data in the records. The source activity at each region is represented as a row in a matrix, in the same way that MEG sensors are represented in the sensor-level CNN. At the sensor level there is a rough spatial correlation between row index and sensor position, such that adjacent rows in the record tend to represent data from nearby sensors. This spatial correlation is much poorer in our source-estimated representation when reducing 3-dimensional positions to rows in the record. Likely, the associated spatial discontinuities are problematic for training the kernels. In order to fully exploit the advantages of source space estimates and the ability to extract features using convolutions, a different representation is required. This representation could include three dimensions to represent the location of activity in the brain, with temporal dynamics over the fourth dimension. This four-dimensional representation of the data would be much more computationally intensive, as it would mean using 10,000’s of vertices. With sufficient time and GPU access, a 4-D CNN would be an interesting future direction for this study, which may improve classification accuracy on source estimated data.

One main advantage of using deep learning is that feature engineering is not required in order to develop viable and effective models. Guided by probability, neural networks learn representations of the data automatically in order to accomplish a task. Feature engineering employed in more traditional machine learning applications, on the other hand, requires expert-level knowledge of a problem area and model predictions can be sensitive to the choice of representation. Furthermore, these representations may fail to capture the more subtle factors of variability within the data that is required to effectively represent class membership. The representations generated by a deep learner empower models that are both more robust and allow us to gain insight into highly complex systems. Our visualisation results provide evidence that a CNN is effective at identifying features of the neuromagnetic recordings that are previously known to be generated during cued button pressing. This is an important validation of the proposed CNN before its utilisation in more challenging problems (i.e., more complicated cognitive tasks).

Saliency maps appeared to provide more consistent results than the occlusion histograms, which showed a more uniform distribution over the baseline records. This intuitively suggests that specific (occluded) regions were more important for classifying active records as compared to the baseline ones. Visual inspection of the topographic plots showed that the saliency maps tended to more accurately capture the regions of activation as demonstrated within the evoked field. Furthermore, the late component represented in the saliency maps (approximately 200-400 ms) may represent a sensitivity to suppression of cortical rhythms commonly observed in movement tasks.

Further work is required in order to use CNNs applied to MEG data in a clinical setting. Ideally, a system employing these networks could provide an informative model of healthy and pathological brain activity for a given paradigm. Such a network could provide classification for diagnosis or treatment planning, including visualisation to localise regions of the brain presenting the activity of interest. Our work suggests that large datasets of neuromagnetic recordings will be required to make this clinical application a reality.

## 5. Acknowledgments

This research was funded by the Natural Sciences and Engineering Council of Canada (grant RGPIN-2018-05470). Data collection and sharing for this project was provided by the Cambridge Centre for Ageing and Neuroscience (CamCAN). CamCAN funding was provided by the UK Biotechnology and Biological Sciences Research Council (grant number BB/H008217/1), together with support from the UK Medical Research Council and University of Cambridge, UK.

